# Arrested PPTC7 import couples mitochondrial depolarisation to NIX and BNIP3 degradation

**DOI:** 10.64898/2026.10.16.759620

**Authors:** Ioanna Ch. Georgiou, Anne Clancy, Emma V. Rusilowicz-Jones, Sylvie Urbé, Michael J. Clague

## Abstract

Mitophagy is initiated by two distinct pathways that are both coupled to mitochondrial protein import. Import stress stabilises PINK1 at the outer mitochondrial membrane, activating the Parkin and ubiquitin-dependent pathway. Receptor-mediated pathways are governed by the expression of outer mitochondrial membrane proteins NIX and BNIP3. Mitophagy is limited by their degradation, via SCF^FBXL4^. This requires the PP2C-family phosphatase PPTC7 as an obligate substrate recognition factor, captured at the outer membrane en route to the matrix. We show that mitochondrial depolarisation exerts opposing effects on these two pathways. In RPE1 cells, depolarisation drives rapid, ubiquitin and proteasome-dependent loss of NIX and BNIP3 that requires ongoing protein synthesis of PPTC7. Using a doxycycline-mediated pulse-chase assay, we demonstrate directly that depolarisation inhibits import of newly synthesised PPTC7, and that this arrested PPTC7 is itself degraded in an FBXL4-dependent manner alongside its substrates. Thus, import arrest both stabilises PINK1 and eliminates NIX and BNIP3, coordinating a switch between mitophagy pathways, while concurrent degradation of PPTC7 couples mitophagy receptor turnover to PPTC7 synthesis rates.

## Introduction

The selective autophagy of mitochondria, known as mitophagy, maintains mitochondrial quality control and permits remodelling in response to metabolic demand (Clague & Urbe, 2025). The Parkinson’s disease-associated kinase PINK1 acts as a sensor of mitochondrial import stress, notably that arising from mitochondrial depolarisation (Narendra & Youle, 2024). PINK1 accumulation at the outer mitochondrial surface leads to its activation and consequent phosphorylation at Ser65 of ubiquitin conjugated to mitochondrial membrane proteins. This recruits the E3 ligase Parkin, itself activated by direct PINK1-dependent phosphorylation, also on Ser65 (Kane et al, 2014; Kazlauskaite et al, 2014; Kondapalli et al, 2012; Koyano et al, 2014). The resulting accumulation of ubiquitin and phospho-ubiquitin at the outer mitochondrial membrane (OMM) surface recruits mitophagy adaptors, such as optineurin and NDP52. These link to phagophore membranes, by binding to LC3 and other core components of the autophagy machinery, such as TBK1 and FIP200 (Heo et al, 2015; Lazarou et al, 2015; Richter et al, 2016).

A second pathway to mitophagy was originally linked to developmental programmes, such as the co-ordinated removal of mitochondria during reticulocyte and neuronal maturation (Esteban-Martinez et al, 2017; Ordureau et al, 2021; Sandoval et al, 2008; Schweers et al, 2007). This involves the upregulation of the mitophagy receptor BNIP3L, hereafter referred to as NIX. The single-pass transmembrane proteins, NIX, and its closely related paralogue BNIP3, are tail-anchored proteins which directly insert into mitochondrial membranes. From there they can promote mitophagy, independently of ubiquitin, by interaction with LC3 and with the core autophagy machinery components WIPI2/3 (Adriaenssens et al, 2025; Bunker et al, 2023; Hanna et al, 2012; Novak et al, 2010; Zhang et al, 2012). BNIP3 is strongly linked to hypoxia - induced mitophagy and is transcriptionally up-regulated by HIF-1α (Bellot et al, 2009). The Cullin RING ubiquitin ligase 2 (CRL2) substrate receptor VHL limits mitophagy by suppressing HIF-1α-mediated transcription of BNIP3, and to a lesser extent, NIX. Both NIX and BNIP3 are additionally controlled post-translationally by the ubiquitin E3 ligase SCF^FBXL4^, which promotes their degradation (Cao et al, 2023; Elcocks et al, 2023; Nguyen-Dien et al, 2023). FBXL4, is mutated in a mtDNA depletion syndrome, causing severe paediatric disease (Alsina et al, 2020; Bonnen et al, 2013; Chen et al, 2023; Gai et al, 2013).

SCF^FBXL4^ is unconventional in requiring an additional component, the PP2C - family phosphatase PPTC7, for recognition of its substrates BNIP3 and NIX (Nguyen-Dien et al, 2024; Sun et al, 2024; Wei et al, 2024). PPTC7 carries a 13 amino acid “weak” mitochondrial targeting signal that is removed upon import. Accumulation of BNIP3 or NIX on the OMM enables PPTC7 precursor capture prior to import, and retention at the OMM. Here it assembles with the SCF^FBXL4^ complex resulting in BNIP3/NIX degradation. The size of the OMM-PPTC7 pool is set by the balance between import and capture. Thus, OMM-PPTC7 acts as a rate-limiting rheostat for BNIP3 and NIX-dependent pathways (Nguyen-Dien et al., 2024; Sun et al., 2024; Wei et al., 2024).

In this manuscript we consolidate a model for a co-ordinated switch between these mitophagy pathways; whilst PINK1 accumulates, NIX and BNIP3 are actively degraded. Using a combination of knock-out cell lines and inducible expression we show that nascent PPTC7 import is arrested upon mitochondrial depolarisation leading to FBXL4-dependent degradation of NIX and BNIP3. We also reveal that under these conditions, OMM-PPTC7 fails to accumulate, being itself degraded, in an FBXL4 and proteasome-dependent manner. The concurrent degradation of an obligatory substrate recognition factor and substrates provides a means of coupling this degradation pathway to PPTC7 synthesis rates.

## Results

### Mitochondrial depolarisation leads to NIX and BNIP3 degradation

We used a combination of respiratory chain inhibitors, Antimycin and Oligomycin (AO) to induce mitochondrial depolarisation in RPE1 (Flp-In) cells, which express PINK1 but lack the signal amplifier Parkin. This leads to PINK1 and pSer65-ubiquitin accumulation, which is concurrent with a striking depletion of both mitophagy receptors BNIP3 and NIX, which reaches ∼40% and 60% respectively after 4 hours (Figure 1A, B). Other mitochondrial markers that we have tested remain stable at that time point (Figure 1C). Although, AO-dependent NIX and BNIP3 loss is coincident with PINK1 accumulation, there is no contingency on PINK1 availability as their decay is equally evident in PINK1 KO cells (Supplementary Figure S1). Note that under our experimental conditions we do not detect any SDS-resistant dimeric forms of NIX or BNIP3. To summarise, depolarisation induces an increase in PINK1/ubiquitin-dependent mitophagy signals whilst simultaneously repressing the NIX and BNIP3 mitophagy receptors (Figure 1D).

**Figure 1.**
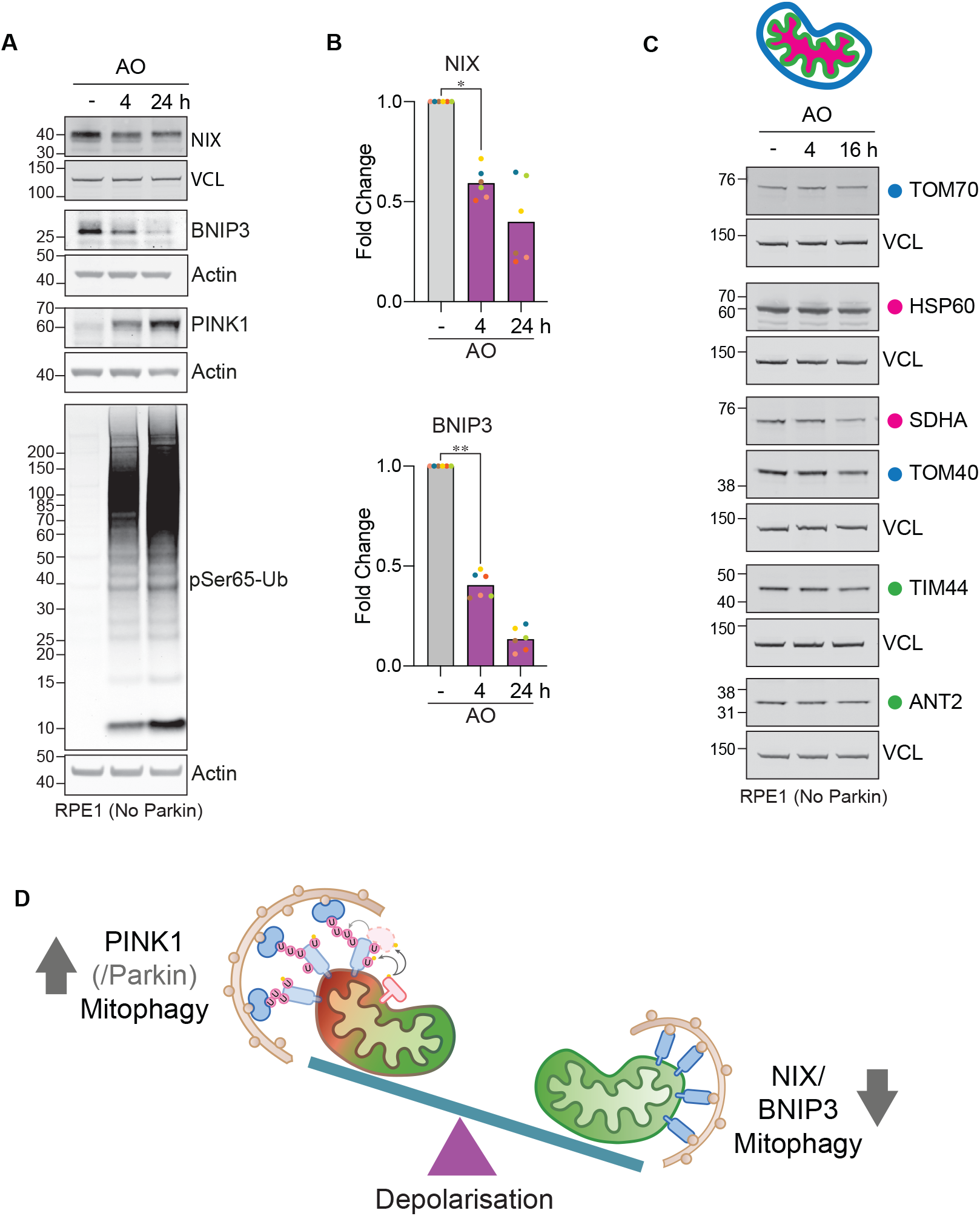
Depolarisation selectively triggers loss of NIX and BNIP3. **A**. RPE1 (Flp-In) cells were treated with or without Antimycin and Oligomycin (AO, 1 and 10 *µ*M respectively) for the indicated times. **B**. Quantification of NIX and BNIP3 levels for six independent, colour-coded experiments represented in A. Paired t-test were performed on vehicle control (-) versus 4 h with data normalised to the 24 h sample to account for gel-to-gel signal variability. NIX *p=0.0110, BNIP3 **p=0.0068. **C**. Survey of mitochondrial proteins following depolarisation. RPE1 (Flp-In) cells were treated with AO for the indicated times. Coloured dots indicate OMM (blue), IMM (green) and Matrix or Matrix-facing (pink) proteins. **D**. Schematic illustration of the depolarisation-induced mitophagy pathway switch. VCL; Vinculin.

We asked if this apparent degradation of BNIP3 and NIX is ubiquitin-dependent by adding the ubiquitin E1 activating enzyme inhibitor, TAK-243 (±AO). We observed TAK-243 blocks AO-dependent degradation of NIX and BNIP3 (Figure 2A,B). Depending on the exact running conditions of SDS-PAGE, NIX can be resolved as a doublet, where the upper band represents a phosphorylated species (Figure 2C). In order to overcome any confounding effects of differential antibody recognition of proteoforms, we have treated samples with λ-phosphatase to collapse all species back to an unphosphorylated form. For the set of antibodies used in these experiments, the recovered intensity of unphosphorylated NIX is equal to that derived from a mixture of proteoforms. BNIP3 differs, in that the measured intensity is increased upon phosphatase treatment, but the fold decrease in response to AO is maintained (Figure 2C,D). We conclude that the apparent loss of NIX and BNIP3 does not represent conversion between phosphorylation states. Concordantly, we have re-expressed epitope-tagged HA-NIX in BNIP3/NIX RPE1 (Flp-In) double KO cells, for which we also observe an AO-dependent enhancement of its turnover (Figure 2E,F).

**Figure 2.**
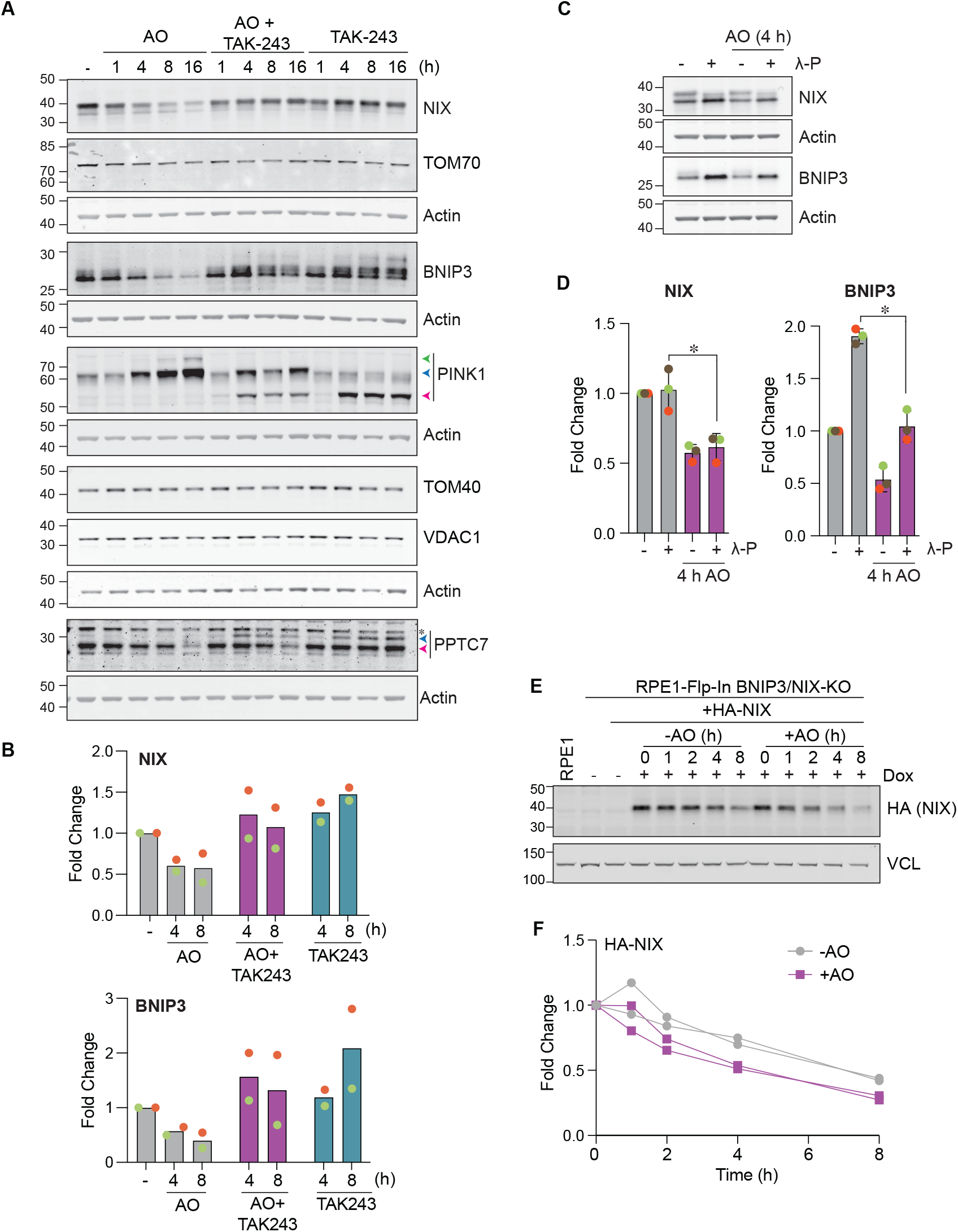
Depolarisation-induced NIX/BNIP3 degradation is ubiquitin-dependent. **A**. RPE1 (Flp-In) cells were treated with Antimycin and Oligomycin (AO, 1 and 10 *µ*M respectively) and/or TAK-243 (1 *µ*M) for the indicated times. Blue and pink arrowheads indicate precursor and matrix forms of PPTC7; green, blue and pink arrowheads indicate mono-ubiquitylated, full-length and cleaved species of PINK1, respectively. * indicates non-specific band **B**. Quantification of two independent, colour-coded experiments represented in A. Data are displayed as fold change relative to vehicle control (-). **C**. RPE1 (Flp-In) cells were incubated with or without AO for 4 h and lysates treated with or without λ-Phosphatase (λ-P). **D**. Quantification of three independent experiments represented in C. Data are displayed as fold change relative to vehicle control (-). Error bars indicate standard deviation. NIX *p-value=0.0117, BNIP3 *p-value=0.0142. **E**. RPE1 BNIP3/NIX KΟ (Flp-In) cells with stably integrated HA-NIX were treated with or without Doxycycline (Dox, 10 ng/ml) for 6 h to induce expression, then chased in fresh media (without Dox) for 16 h before AO-treatment for the indicated times. VCL; Vinculin. **F**. Quantification of two independent experiments represented in E. Data are displayed as fold change relative to t=0.

We used a suite of reagents to interrogate both the basal turnover and the AO-induced degradation pathway of NIX and BNIP3. In addition to TAK-243, AO-induced NIX degradation is rescued by the proteasomal inhibitors MG132 and epoxomicin and to a much lesser degree by lysosomal v-ATPase inhibitors concanamycin A and bafilomycin A (Figure 3A,B). Under basal conditions, BNIP3 levels clearly increase with both proteasomal and lysosomal inhibitors, consistent with previous findings (Figure 3A,C) (Delgado et al, 2024). However, the lysosomal degradation pathway is completely overridden under depolarisation conditions, wherein the relevant inhibitors (Baf A, Conc A) provide no rescue and both NIX- and BNIP3-degradation are solely sensitive to proteasomal inhibition. Note that both NIX and BNIP3 accrue to the same extent with proteasomal inhibitors, under basal and AO-treated conditions. MLN-4924, which inhibits all CRL E3-ligase activity by blocking Cullin neddylation, elevates both NIX and BNIP3 (Figure 3A-C), through a combined effect on VHL (transcriptional regulation) and FBXL4 (increased stability) (Elcocks et al., 2023; Soucy et al, 2009). We also looked at PPTC7 under these same conditions. Without inhibitors, only the imported matrix form is clearly evident, but a higher molecular weight precursor form is apparent upon MLN4924, TAK-243 or proteasome inhibitor treatment, both with and without AO treatment.

**Figure 3.**
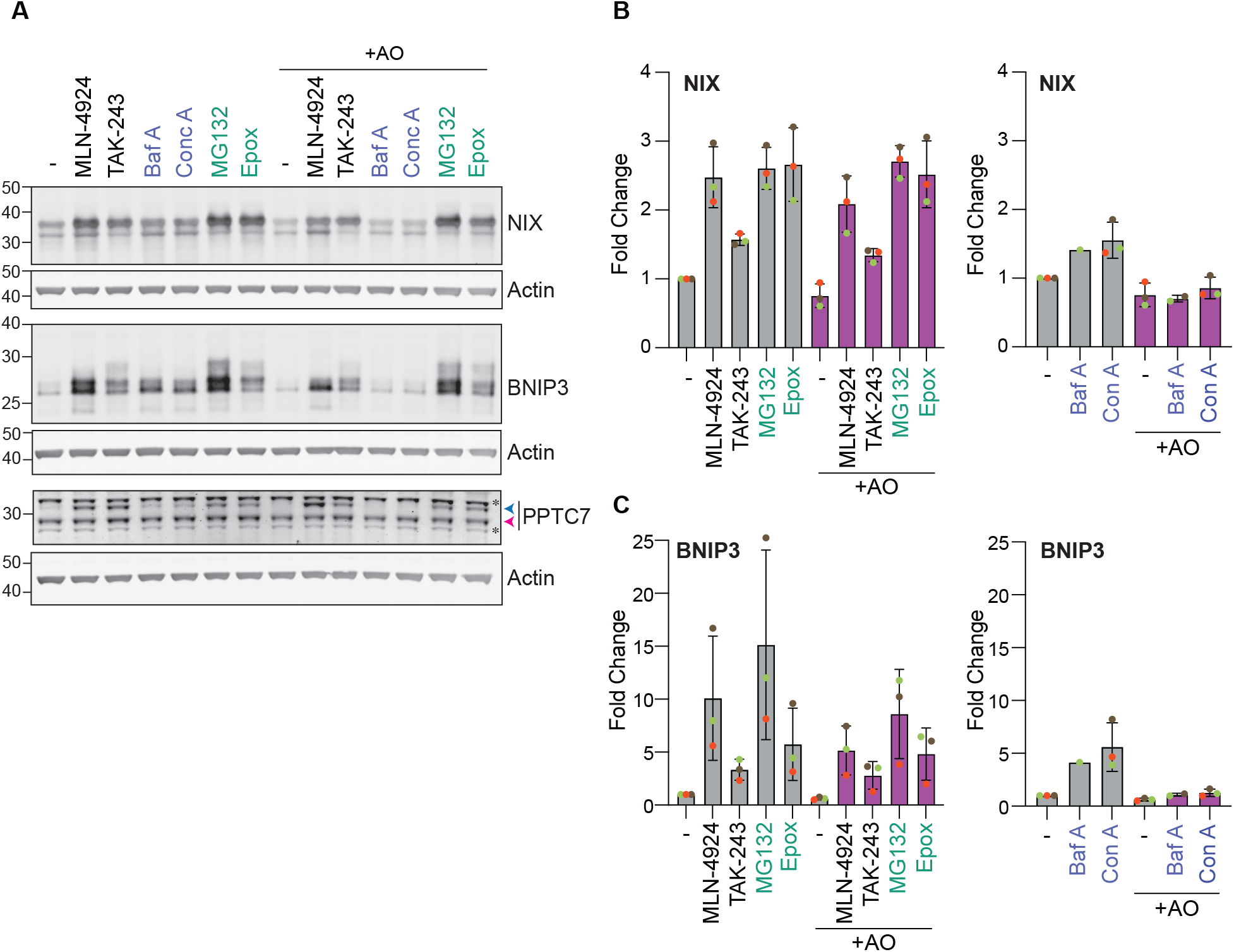
Depolarisation-induced turnover of NIX and BNIP3 are proteasome-dependent. **A**. RPE1 (Flp-In) cells were treated for 8 h with MLN-492 (1 *µ*M), TAK-243 (1 *µ*M), Bafilomycin A1 (Baf A, 100 nM), Concanamycin A (Conc A, 100 nM), MG132 (10 *µ*M) or Epoxomicin (Epox, 10 *µ*M), with or without Antimycin and Oligomycin (AO, 1 and 10 *µ*M respectively). Blue and pink arrowheads indicate PPTC7 precursor and matrix forms respectively. * indicates non-specific bands. **B, C**. Quantification of three independent, colour-coded experiments (except for Baf A), represented in A. Error bars indicate standard deviation. Data are displayed as fold change relative to vehicle treated controls, which are shown in both left and right hand panels.

### AO-**induced BNIP3 and NIX degradation is contingent on FBXL4 and PPTC7 import failure**

NIX and BNIP3 degradation is dependent upon the SCF^FBXL4^-PPTC7 complex (Cao et al., 2023; Chen et al., 2023; Elcocks et al., 2023; Nguyen-Dien et al., 2023; Nguyen-Dien et al., 2024; Sun et al., 2024; Wei et al., 2024). We have generated isogenic RPE1 (Flp-In) KO cell lines for FBXL4 and PPTC7 as well as a double KO line for BNIP3 and NIX. Both FBXL4 and PPTC7 KO cells show elevated BNIP3 and NIX levels whilst other mitochondrial markers tend to decrease in line with previous findings (Figure 4A-F) (Alsina et al., 2020; Niemi et al, 2019). In PPTC7 and FBXL4 KO cell lines, the early (i.e. 4 hours) NIX/BNIP3 degradative response to AO is lost (Figure 4A-C and Supplementary Figure S2A-C). Treating samples with λ-phosphatase to collapse phosphorylation reinforces this finding (Figure 4H,I). We do however note a higher molecular weight λ-phosphatase resistant form of NIX in FBXL4 and PPTC7 KO cells. Collectively our data verify that the FBXL4/PPTC7 pathway is operative in AO-induced degradation of NIX and BNIP3 and demonstrate that PPTC7 remains an essential factor.

**Figure 4.**
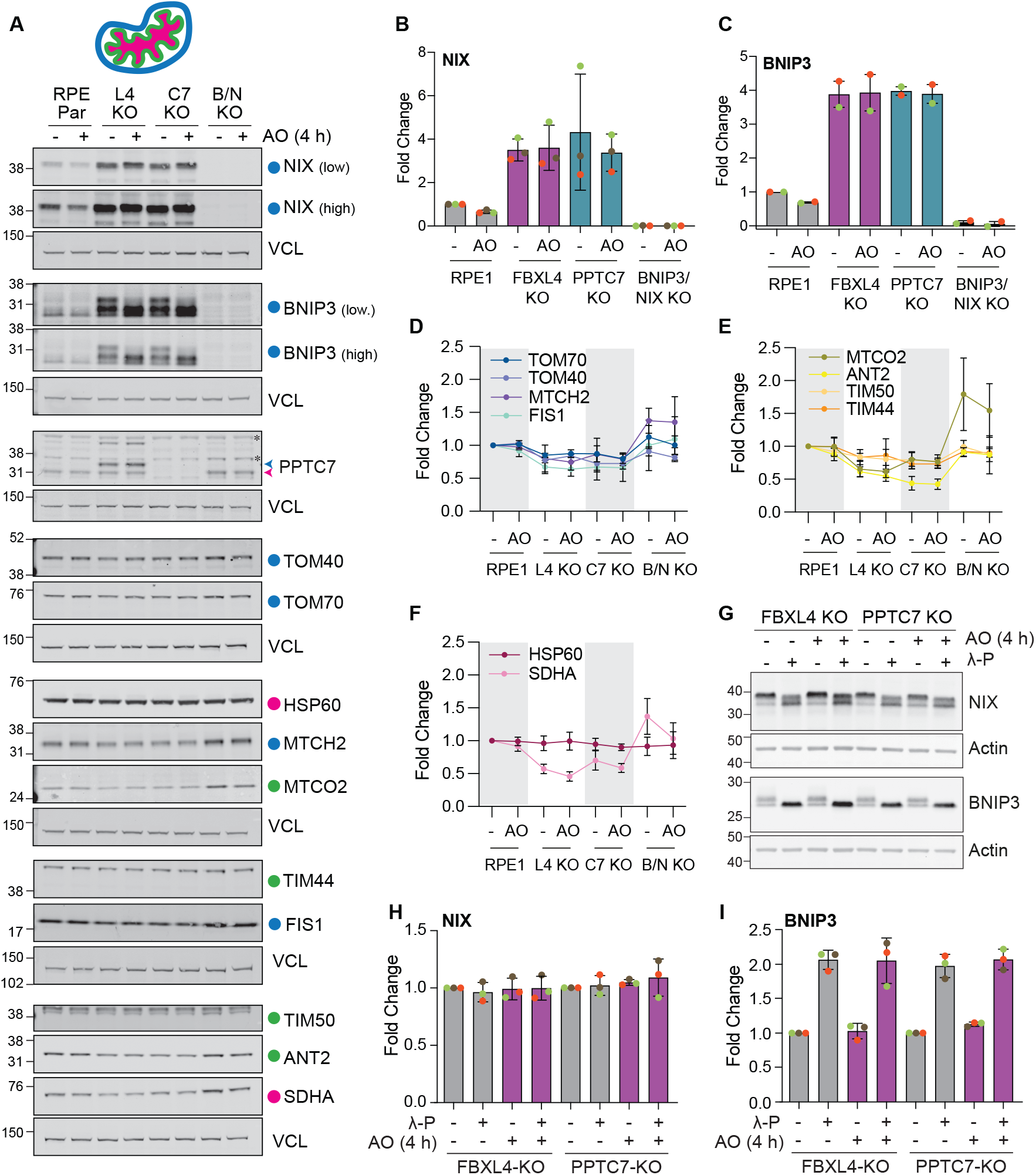
Depolarisation-induced NIX and BNIP3 decay is suppressed in FBXL4 and PPTC7 KO cells. **A**. Parental (RPE Par), FBXL4 (L4), PPTC7 (C7) and BNIP3/NIX (B/N) KO RPE1 (Flp-In) cells were treated for 4 h with or without Antimycin and Oligomycin (AO, 1 and 10 *µ*M respectively). Coloured dots indicate OMM (blue), IMM (green) and Matrix and matrix-facing (pink) proteins. Blue and pink arrowheads indicate PPTC7 precursor and matrix forms respectively. VCL; Vinculin, * indicates non-specific bands. **B, C**. Quantification of three (NIX) or two (BNIP3) independent, colour-coded experiments represented in A. Data are displayed as fold change relative to vehicle treated. Error bars indicate standard deviation (NIX) or range (BNIP3). **D**-**F**. Quantification of OMM, IMM and Matrix/matrix-facing mitochondrial proteins represented in A. Error bars indicate standard deviation, n=3. **G**. FBXL4 and PPTC7 KO RPE1 (Flp-In) cells were treated as in A and lysates treated with or without λ-Phosphatase (λ-P). **H**,**I**. Quantification of three independent, colour-coded experiments, represented in G. Error bars indicate standard deviation. Data are displayed as fold change relative to vehicle treated.

For our model to hold true, depolarisation must inhibit the import of PPTC7, but this has not so far been formally shown. We do not see any accumulation of a precursor form of PPTC7 (OMM-PPTC7) upon depolarisation (see Figures 2A and 3A). We therefore developed a pulse-chase assay to visualise PPTC7 import, using a pulse of doxycycline-inducible expression of PPTC7-HA in RPE1 (Flp-In) PPTC7 KO cells. Cells are treated with doxycycline (1 hour), which is then washed out and chased with fresh medium for indicated times (Figure 5A). The kinetics are such that we can clearly capture the OMM form of PPTC7-HA and follow its processing to the mature matrix form (Figure 5B,C, blue and pink arrowheads respectively). Doxycycline induction of PPTC7-HA expression leads to loss of NIX and BNIP3, which later rebounds just as levels of the PPTC7 precursor form decline (Figure 5D,E). Neither import rate nor NIX/BNIP3 degradation are influenced by D290N mutation in PPTC7 which has previously been shown to largely inactivate phosphatase activity (Figure 5B, Supplementary Figure 3A) (Nguyen-Dien et al., 2024). Importantly, mitochondrial import of PPTC7-HA is completely blocked by AO treatment, both in reconstituted PPTC7 KO cells, and in a RPE1 (Flp-In) parental cell background (Figure 5 F, G, H and Supplementary Figure S3 B, C). However, under these conditions, OMM-PPTC7 levels still decline over the chase period reflecting an otherwise cryptic degradation pathway. The early component of this degradation (up to 4 h) is absent in FBXL4 KO cells (Figure 5I, J). At a later time point (24 h), an alternative degradation pathway has cleared remaining OMM-PPTC7 independently of FBXL4 status.

**Figure 5.**
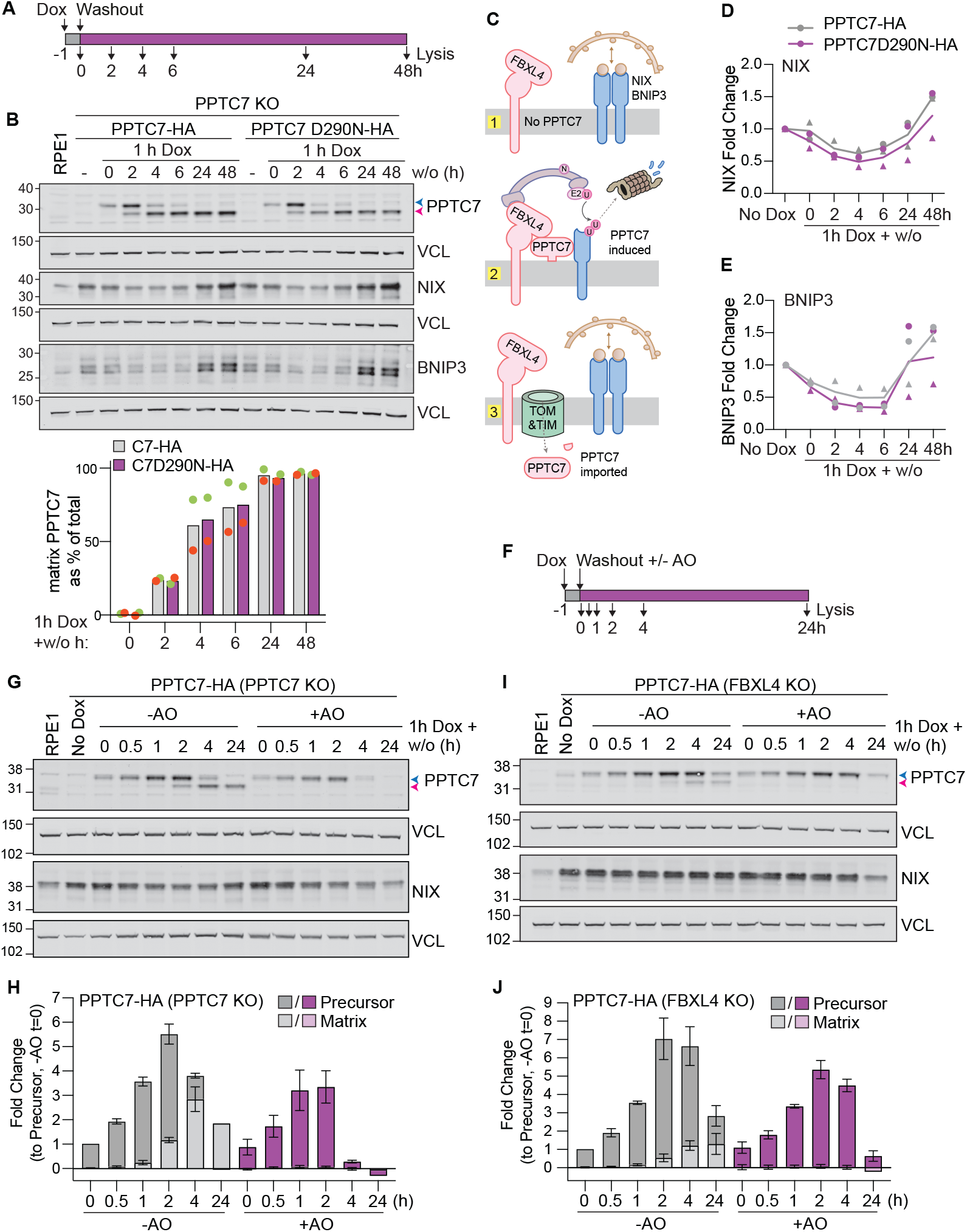
PPTC7 import is blocked upon depolarisation and its precursor is degraded by FBXL4. **A, B**. RPE1 PPTC7 KO (Flp-In) cells with stably integrated wild-type PPTC7-HA or a phosphatase-deficient PPTC7-D290N-HA mutant, were treated with Doxycycline (Dox, 10 ng/ml) for 1 h to induce expression, then incubated in fresh media without Dox for indicated times (w/o; wash-out). Representative western blot (n=2) and associated quantification of PPTC7 import showing matrix PPTC7 as % of total PPTC7. Blue and pink arrowheads indicate PPTC7 precursor and matrix forms respectively. **C**. Schematic illustrating the PPTC7 status and impact on NIX for for the following conditions: (1) No dox (-); (2,) 2 h w/o; (3) 24 h w/o. **D, E**. Quantification of NIX (D) and BNIP3 (E) protein levels for two independent experiments represented in B. **F**. Schematic illustrating the sequential treatments for experiments shown in G - J. **G**. RPE1 PPTC7 KO (Flp-In) cells with stably integrated PPTC7-HA were treated with Doxycycline (Dox, 10 ng/ml) for 1 h to induce expression, then incubated in fresh media (no Dox) and treated with or without Antimycin and Oligomycin (AO, 1 and 10 *µ*M respectively) for the indicated times, (w/o; wash-out). **H**. Quantification of experiments represented in G. Data shown are normalised to the precursor form of PPTC7 after 1 h Dox induction (t=0). Error bars indicate standard deviation (n=3; except 24 h timepoint, n=1). **I**. RPE1 FBXL4 KO (Flp-In) cells with stably integrated PPTC7-HA were treated with Doxycycline (Dox, 10 ng/ml) for 1 h to induce expression, then incubated in fresh media (no Dox) and treated with or without AO for the indicated times. **J**. Quantification of two independent experiments represented in I. Data shown are normalised to the precursor form of PPTC7 after 1 h Dox induction (t=0, -AO). Error bars show range. VCL; Vinculin.

## Discussion

Here we explore the concept of PPTC7 as a mitochondrial import stress sensor. We have used RPE1 (Flp-In) cells as an experimental platform. They are an established model for FBXL4 regulation of BNIP3 and NIX and allow the comparison of wild-type and mutant proteins in an isogenic setting. They also lack Parkin, ensuring a more stable mitochondrial population in the face of depolarisation, thereby removing a factor that could complicate our analysis.

The same conditions that turn the PINK1/ Parkin pathway on via PINK1 accrual, also lead to reductions in mitophagy receptors BNIP3 and NIX, through the arrest of PPTC7 mitochondrial import. Thus, we propose a depolarisation-induced switch between pathways, that is enabled by the combined effect of two import-arrest sensors. In this framing, the respective mitophagy arms are not redundant back-ups but contextually prioritised. Receptor-mediated mitophagy covers basal, programmed and hypoxic turnover whilst acute damage triggers the PINK1/Parkin pathway and actively dismantles the receptor route.

PPTC7 suppresses mitophagy by connecting the SCF^FBXL4^ complex to its substrates, NIX and BNIP3 (Nguyen-Dien et al., 2024; Sun et al., 2024; Wei et al., 2024). The PPTC7 phosphatase fold in this context acts as a substrate recognition module, since a catalytic inactivating mutation (D290N) has no effect on either import or receptor degradation kinetics (Figure 5A-D) depolarisation on NIX and BNIP3 is primarily at the level of the mitochondrial import block to PPTC7 rather than translation. NIX and BNIP3 accumulate to the same levels following proteasome inhibition under AO and basal conditions (Figure 3A,B) and PPTC7 generation is itself largely insensitive to AO (Figure 5).

In contrast to PINK1, we do not see any accrual of PPTC7 at the OMM following depolarisation. This is because it too is efficiently degraded in an FBXL4-dependent manner (Figure 5G,I). Note that, these mechanistic findings reflect the early response (≤ 4 hours) to depolarisation, whereas later phases may reflect more general proteostatic events, including mitophagy itself. The concurrent degradation of adaptor and mitophagy receptor ensures that the extent of receptor loss reflects PPTC7 biosynthesis under conditions of import stress as well as under basal conditions.

Our study introduces an induced PPTC7 expression and pulse-chase protocol, as a means to gain insight into its mitochondrial import and its role in the FBXL4-mediated degradation pathway. We discovered an import stress-dependent loss of the BNIP3/NIX mitophagy receptors, coincident with PINK1/Parkin pathway activation, which may have physiological significance. FBXL4 provides a brake on non-selective mitophagy and its loss-of-function results in encephalomyopathic mtDNA depletion syndrome (Alsina et al., 2020; Bonnen et al., 2013; Gai et al., 2013). OMM-PPTC7 effectively determines how firmly the brake can be applied and is made more available under conditions of import failure, whilst the PINK1 pathway is up-regulated. An analogy can be made with prioritisation of DNA double-strand break repair processes. Cells can choose between accuracy and fidelity by glueing the ends straight back together (non-homologous end joining, NHEJ) or rebuilding the break using an intact copy as a template (homologous recombination). Switching to homologous recombination requires actively removing Ku70/80, which otherwise commits the ends to NHEJ (Chapman et al, 2012).

The PINK1-mediated damage response is multilayered and thresholded; a lag time precedes commitment to mitophagy following acute depolarisation and ubiquitin-dependent mitophagy signals may be opposed by deubiquitylases such as USP30 (Bingol & Sheng, 2016; Liang et al, 2015; Pollock et al, 2021). Suppressing non-selective mitochondrial turnover may provide a means to preserve the highest possible pool of mitochondria whilst triage towards the PINK1/Parkin pathway is occurring. In summary, we propose that mitochondrial import competence is sensed by two short-lived mitochondrial proteins, PINK1 and PPTC7. One licenses ubiquitin-dependent mitophagy and the other withdraws the receptor-mediated route. Rather than acting as a simple trigger, depolarisation reconfigures which arm of mitophagy operates.

## Acknowledgements

This work was supported by the Medical Research Council MR/Y012356/1. MJC was recipient of a Royal Society industry fellowship, INF\R2\212031. We thank Sanchitha Shetty Rajkumar for her help with preliminary experiments.

## Methods

### Cell culture

hTERT-RPE1 Flp-In T-REx cells (a gift from Jonathon Pines, London, UK), referred to as RPE1 (Flp-In), and engineered lines described below, were cultured in Dulbecco’s modified Eagle’s medium DMEM-F12 supplemented with 10% FBS.

### Generation of knockout (KO) cell lines

PINK1 Knockout cells were described in Pollock et al. (Pollock et al, 2025). To generate CRISPR/Cas9 knockout (KO) lines, annealed sgRNAs were cloned into pU6-(BbsI)-CBh-Cas9-T2A-mCherry (Addgene, #64324), transiently transfected into hTERT-RPE1 Flp-In T-REx cells. Twenty-four hours later, mCherry-positive cells were selected by fluorescence-activated cell sorting (FACS). Individual clones were isolated by single cell dilution and validated by western blotting. Guide sequences w e r e a s f o l l o w s : B N I P 3 : 5 ‘ - GAGAGAAAAACAGCTCAC-3’, FBXL4 : 5’-CTCAATGCAGAGGTAGTCCA-3’, NIX: 5’-TAGCTCTCAGGTGTGTCGGG-3’, PPTC7: 5’-TTCGTACCTAGTAATCCCAT-3’.

### Generation of RPE1 Flp-In T-REx inducible expression cell lines

HA-NIX and PPTC7-HA constructs were cloned into pcDNA5 FRT-TO NeoR or pcDNA5-FRT-TO PuroR, respectively, by Gibson assembly and transfected into hTERT-RPE1 Flp-In T-REx alongside pOG44 (1:9) using Lipofectamine 3000. Forty-eight or twenty four hours after transfection, positive cells were selected with Blasticidin (10μg/ml) and G418 (400μg/ml) or Puromycin (1 µg/ml). Cell pools were validated by western blotting. Expression of transgenes was induced with 10 ng/ml Doxycycline for the indicated time.

### Antibodies and other reagents

Antibodies are as follows: anti-BNIP3 (rabbit, 1 : 1000, ab109362 Abcam) , anti - BNIP3 (rabbit,1:1000, 44060, Cell Signaling Technology), anti-BNIP3L/NIX (rabbit, 1:1000-1:2000, 12396, Cell Signaling Technology), anti-PPTC7 (rabbit, 1:1000-1:2000, NBP1-90654, Novus), anti-PINK1 D8G3 (rabbit, 6946, 1:1000, Cell Signaling Technology), pSer65Ub (rabbit, 1:1000, 62802,Cell Signaling Technology), anti-Actin (mouse, 66009, 1:20,000, Proteintech), anti-vinculin (mouse, 1:1000-1:2000, Santa Cruz sc-55465), anti-HA (mouse, 1:1000, Biolegend MMS-101P) anti-TOMM20 (rabbit, 11802-1-AP, 1:1000, Proteintech), TOMM70 (mouse, 1:1000, SantaCruz sc-390545), TOMM40 (rabbit 1:1000, Novus NBP2-38289), TIMM44 (rabbit, 1:1000, Sigma HPA043052), TIMM50 (C-9, mouse, 1:1000, Santa Cruz sc-393678), Fis1 (rabbit, 1:1000, Protein Tech 10956-1-AP), ANT2 (rabbit, 1:1000, CST 14671), HSP60 (N-20, Goat, 1:1000, SantaCruz sc-1052), SDHA (rabbit 1:1000, CST 11998), VDAC1 (mouse 1:1000, Abcam ab14734), MTCO2 (12C4F12, mouse 1:1000, Abcam ab110258), MTCH2 (rabbit, 1:1000, Abcam ab113707).

Other reagents: oligomycin A (75351, Sigma-Aldrich), antimycin A (A8674, Sigma-Aldrich), doxycycline (D9891, Sigma-Aldrich), epoxomicin (324800, Sigma-Aldrich), TAK-243 (MLN7243, S8341, Selleckchem), bafilomycin A (B1793, Sigma-Aldrich), concanamycin A (C9705, Sigma-Aldrich), MG132 (474790, Sigma-Aldrich), MLN-4924 (Pevonedistat, C-1231, Chemgood), Lambda Protein Phosphatase (P0753, New England Biolabs).

### Cell lysate preparation

Cells were lysed in NP-40 lysis buffer (0.5% Nonidet P-40 substitute (74385 Sigma), 25 mM Tris–HCl, pH 7.5, 100 mM NaCl, and 50 mM NaF) supplemented with mammalian protease inhibitors (MPI, Sigma-Aldrich) and PhosSTOP (Roche).

### Western blotting

Proteins were resolved using 4-12% NuPage gels (Invitrogen), transferred to nitrocellulose membranes and blocked in either 5% milk, or 5% fat-free bovine serum albumin in tris-buffered saline (TBS) with 0.1% Tween20, before primary antibody incubation in the same blocking solution. Western blot visualisation was performed using IRdye 800CW and 680LT coupled secondary antibodies with an Odyssey infrared scanner (Li-COR, Biosciences). For quantification, signal values were obtained from Image Studio software after subtracting the background signal. In all figures, the signal for the protein of interest was normalised to loading control and displayed as a fold change versus the relevant control. For graphs shown in Figures 5H, 5I, S3A and S3C, PPTC7 background values from the un-induced sample (-Dox) were subtracted first before calculating fold change.

### λ-Phosphatase treatments

Cells were treated as indicated before lysis in phosphatase compatible lysis buffer (20 mM Tris-HCl, pH 7.5, 150 mM NaCl, 1 mM EDTA, 1 mM EGTA, and 1% Triton X-100), supplemented with MPIs. 20 μg from each lysate were mixed with 5 μl of 10x NEBuffer for protein metallo-phosphatases (PMP), 5 μl of 10 mM MnCl2, and with or without 1.5 μl of Lambda (λ) protein phosphatase. Reactions were incubated at 30ºC for 1.5 h in a PCR cycler. 12.5 μl of 5x sample buffer was added to each reaction and samples heated for 5 min at 95ºC.

### Statistics

All statistical analysis was done using Prism 11. In figure 1B, vehicle and 4 h time point samples were normalised to the 24 h sample to account for gel-to-gel variability and differing loading controls. Two-way paired T-tests were performed. For Figure 2D, data were first normalised to untreated control (-AO, -λ-P) and a two-way paired T-test was applied to compare λ-phosphatase treated samples ±AO.

## Supplementary Figures

**Figure S1.**
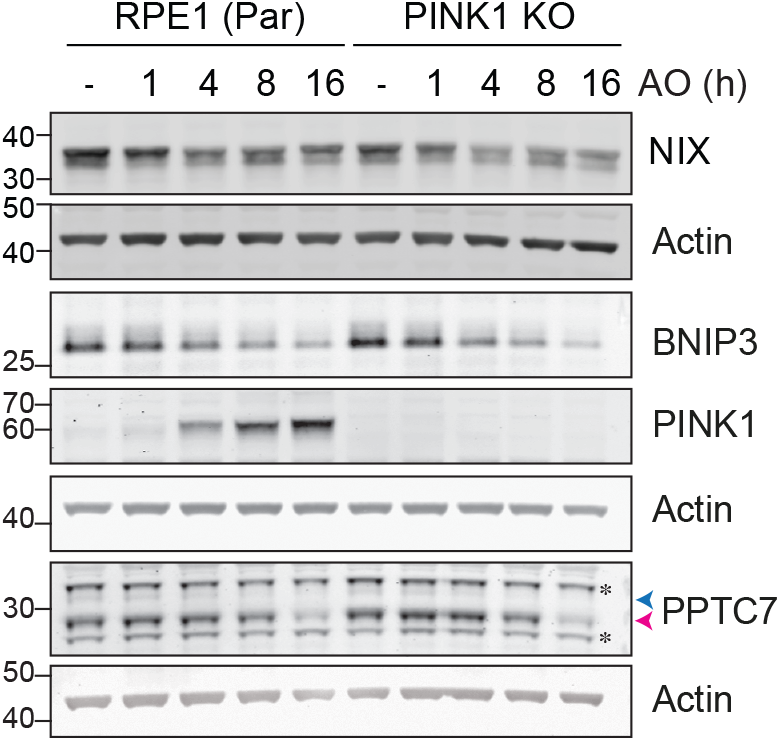
Depolarisation-induced NIX and BNIP3 loss is PINK1-independent. Parental and PINK1 KO RPE1 (Flp-In) cells were treated with or without Antimycin and Oligomycin (AO, 1 and 10 *µ*M respectively) for the indicated times. Blue and pink arrowheads indicate PPTC7 precursor and matrix forms respectively, * indicates non-specific bands.

**Figure S2.**
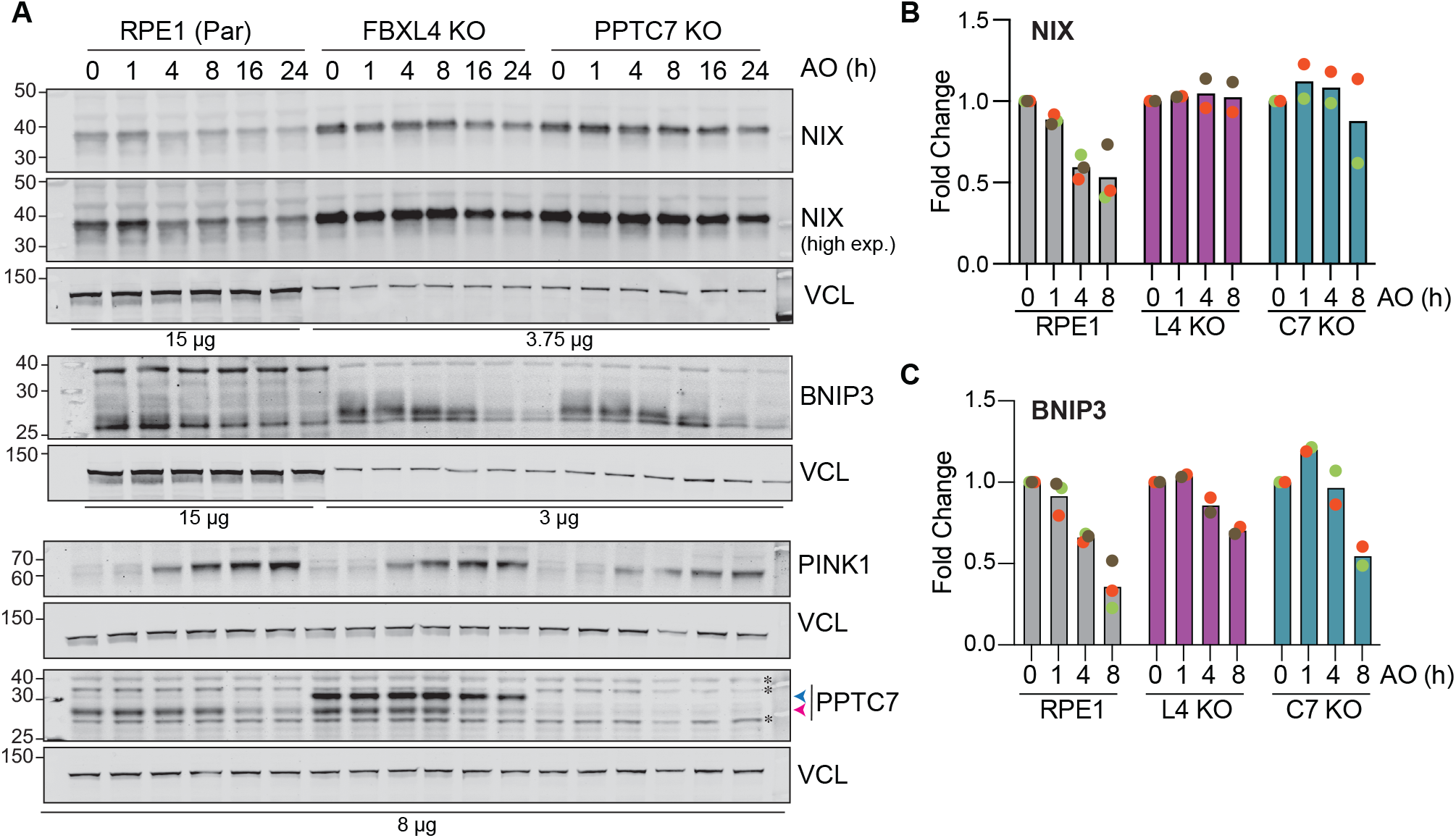
Depolarisation-induced NIX and BNIP3 loss is suppressed in FBXL4 and PPTC7 KO cells(Supplementary to Figure 4) **A**. Parental (Par), FBXL4 KO (L4 KO) and PPTC7 KO (C7 KO) RPE1 (Flp-In) cells were treated with or without Antimycin and Oligomycin (AO, 1 and 10 *µ*M respectively) for the indicated times. Protein loading was adjusted as indicated to allow for a better comparison of NIX and BNIP3 signals, taking into account their increased levels in untreated FBXL4 and PPTC7 KO cells. Blue and pink arrowheads indicate PPTC7 precursor and matrix forms respectively. VCL; Vinculin. **B, C**. Quantification of three (RPE1 Par) two (L4 KO) or two (C7 KO) independent experiments represented in A. Data are displayed as fold change relative to t=0.

**Figure S3.**
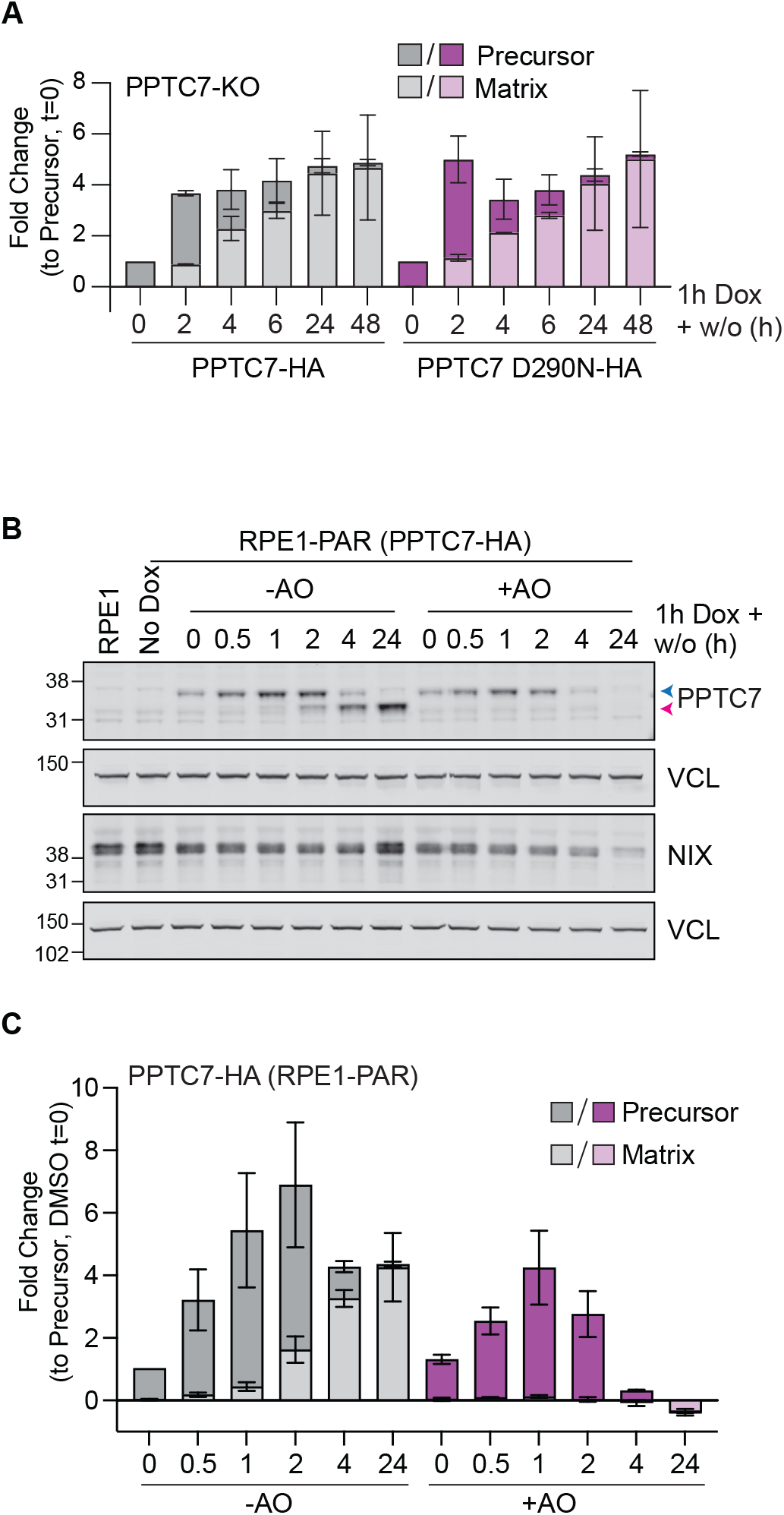
PPTC7 import is sensitive to depolarisation but not affected by phosphatase impairment. (Supplementary to Figure 5) **A**. Quantification of two independent experiments represented in Figure 5B. Data shown are normalised for each cell line to the precursor form of PPTC7 after 1 h Dox induction (t=0). Error bars show range. **B**. RPE1 (Flp-In) cells stably expressing inducible PPTC7-HA were treated with Doxycycline (Dox, 10 ng/ml) for 1 h, then incubated in fresh media (no Dox) and treated with or without Antimycin and Oligomycin (AO, 1 and 10 *µ*M respectively) for the indicated times, (w/o, wash-out). **C**. Quantification of two independent experiments represented in B. Data shown are normalised to the precursor form of PPTC7 after 1 h Dox induction (t=0, -AO). Error bars show range.

